# PersonaAI: An Interactive Agentic-AI Framework for Autonomous Hypothesis Generation and Validation in Aging

**DOI:** 10.64898/2026.01.16.699755

**Authors:** Byounggook Cho, Gi-Young Lee, Junghyun Jung, Junyeop Kim, GunHo Park, Patrick C. N. Martin, Hyobin Kim, Jeein Oh, Jong-Soo Kim, Jongpil Kim, Tae-Hyung Kim, Kyoung-Jae Won

## Abstract

Elucidating the mechanisms of aging is impeded by its stochastic, multi-scale nature and cellular heterogeneity, challenges that are compounded by the overwhelming volume of biomedical literature and the complexity of genome-wide datasets. To overcome these barriers, we present PersonaAI, an interactive agentic-AI framework that acts as a digital co-scientist. By integrating literature-based reasoning with autonomous *in silico* validation, PersonaAI synthesizes over 560,000 aging-related publications via retrieval-augmented generation (RAG) to propose mechanistic hypotheses. These hypotheses are subsequently validated by autonomous agents utilizing single-cell RNA-seq data.

Using a temporal cutoff strategy restricted to pre-2020 literature, we demonstrate that PersonaAI can generate hypotheses effectively validated by post-2021 discoveries, proving its capacity for inference beyond simple information retrieval. In application, the system identified senescent Cirbp+ hepatocytes as a liver-intrinsic aging program and uncovered a middle-aged, male-specific decline in adipose stem and progenitor cells, driven by vascular niche deterioration and disrupted VEGF–VEGFR signaling. These results establish PersonaAI as a scalable platform that augments human intuition with autonomous data-driven validation, providing a generalizable platform for accelerating discovery in aging biology.

## Introduction

Aging is a universal, multi-scale process driving the trajectory of life from development to decline (López-Otín et al., 2023). Unlike acute disease states, aging is defined by profound stochasticity and temporal latency. It manifests as an asynchronous accumulation of damage where neighboring cells in identical tissues may exhibit vastly different molecular signatures (Enge et al., 2017; Martinez-Jimenez et al., 2017). Because this process is driven by stochastic entropy rather than deterministic programming, it represents one of the most intractable analytical challenges in modern biology. This complexity is compounded by the vast temporal scale of mammalian aging, which imposes severe logistical barriers. Establishing rigorous causality between early-life molecular events and late-life phenotypes requires tracking trajectories over decades.

To manage this complexity, the field has prepared large-scale datasets using high-throughput technologies. Comprehensive single-cell transcriptomic atlases (Almanzar et al., 2020; Zhang et al., 2025) now provide a foundational landscape of aging signatures across murine and human lifespans, enabling the *in silico* interrogation of temporal heterogeneity. However, these high-dimensional repositories have created a secondary barrier of interpretation. The ability to distinguish biologically meaningful causal patterns from technical artifacts or “passenger” noise now requires a depth of multi-disciplinary expertise that is impossible to scale through human effort alone (Wilkinson et al., 2016).

The engine of scientific discovery is fueled by the recursive synthesis of accumulated knowledge and empirical validation. Traditionally, researchers have relied on their knowledge to generate hypotheses and reason through potential biological mechanisms. However, with over 560,000 articles indexed in PubMed as of 2025 in aging, the information landscape is too vast for any individual researcher to navigate (Bornmann & Mutz, 2014). This challenge is further intensified by the inherent interdisciplinary nature of gerontology, which demands a simultaneous mastery of evolutionary biology, immunology, metabolism, and neuroscience (Campisi et al., 2019).

To overcome the cognitive and bioinformatic bottleneck, we propose PersonaAI. PersonaAI is an agentic-AI framework designed to function as a digital co-pilot (Boiko et al., 2023; Wang et al., 2023) for aging research. Rather than acting as a passive database, PersonaAI engages in an interactive reasoning loop with the researcher to synthesize existing literature into novel, testable hypotheses. Once a biological mechanism is proposed, the system’s orchestrator autonomously constructs and executes dynamic analytical pipelines, rigorously validating the hypothesis *in silico* by querying high-dimensional single-cell aging atlases.

To provide the necessary biological context, PersonaAI is anchored by a comprehensive knowledge base of over 560,000 aging-related publications. Rather than functioning as a standard search engine, the system utilizes a retrieval-augmented generation (RAG) architecture to reconstruct and synthesize information from diverse biomedical resources (Pan et al., 2024). This allows the system to transcend simple keyword retrieval and performs semantic reasoning to bridge between disciplines, identifying indirect causal links. Operating as a true digital co-scientist, PersonaAI engages in an iterative dialectic with the researcher. Through this interactive feedback loop, the agent assists the human partner in refining broad inquiries into concrete, mechanistically sound, and data-testable hypotheses, effectively accelerating the path from literature review to experimental design. Once a hypothesis is crystallized, PersonaAI deploys a specialized fleet of multi-agent subroutines to orchestrate the *in-silico* validation process. This modular architecture allows the system to autonomously requisition high-dimensional data from the Mouse Aging Atlas and execute complex analytical pipelines.

PersonaAI represents a significant evolution beyond pioneering autonomous frameworks such as Virtual Lab and AI Co-scientist (Boiko et al., 2023; Gottweis et al., 2025; Swanson et al., 2025). While these autonomous partners offer speed and scalability, they face a fundamental efficiency bottleneck: they often rely on unconstrained stochastic exploration, generating vast numbers of hypotheses through a computationally expensive guess-and-check approach. Platforms such as AI Co-scientist (M. Bran et al., 2024) do not possess an integral mechanism for empirical validation. In contrast, PersonaAI diverges from the “brute force” paradigm by operating as a strategic co-pilot. By maintaining human intuition within the reasoning loop, the system effectively constrains the search space to biologically plausible mechanisms. Technically, while earlier systems focused on domains with established predictive models, such as protein folding (Jumper et al., 2021; Swanson et al., 2025), PersonaAI incorporated model context protocols (MCPs) to translate unstructured natural language into executable bioinformatic workflows, enabling researchers to perform complex *in silico* validation of aging mechanisms without specialized coding expertise.

The predictive validity of this architecture is substantiated by our retrospective analysis using temporal cut-off, which serves as a blinded benchmark for prospective discovery. PersonaAI newly identified that the male-specific regenerative decline of Adipose Stem and Progenitor Cells (ASPCs) is not merely intrinsic but is actively orchestrated by the deterioration of the vascular endothelial niche.

PersonaAI functions as an active collaborative partner that engages the researcher in a structured reasoning loop. The system guides logical refinement from broad biological inquiries into precise, mechanistically grounded hypotheses and further assessment based on genomic data.

## Result

### Construction of an Interactive Agentic Framework for Aging Hypothesis Generation and Validation

Aging research faces critical challenges driven by biological stochasticity and temporal latency, which obscure causal mechanisms due to cellular heterogeneity, and the data overload from synthesizing from 565,634 publications (PMC: 212,041 and PubMed: 353,593). To address these issues, we developed PersonaAI, a dual-phase agentic framework (Figure 1).

**Figure 1.**
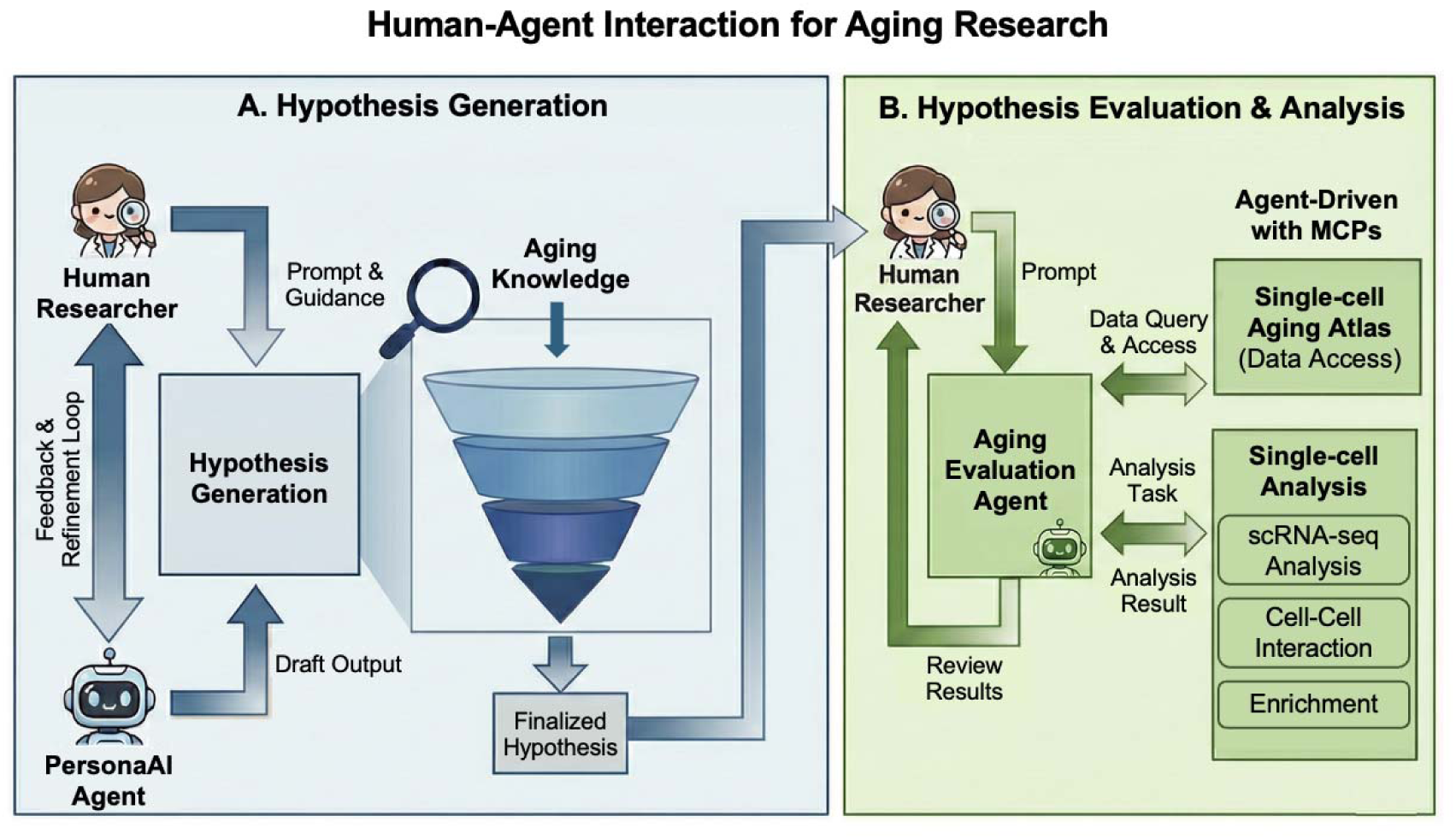
Framework of PersonaAI for autonomous hypothesis generation and in-silico validation in aging research. (A) Schematic of the hypothesis generation phase featuring a Human-Agent interaction loop for literature synthesis and recursive refinement. (B) Evaluation and analysis phase driven by Agent-driven MCPs for autonomous data access and execution of specialized bioinformatic tasks.

Unlike standard Large Language Models (LLMs) that primarily retrieve information, PersonaAI acts as an active research partner through a Human-Agent Interaction Loop (Figure 1A). Rather than acting as a passive knowledge store, the system employs an agentic architecture designed to emulate the iterative nature of scientific reasoning. Through continuous, iterative dialogue, the researcher provides high-level biological guidance, which the system uses to navigate the vast search space of aging literature. This process facilitates knowledge synthesis by bridging disparate datasets and publications to identify hidden biological connections. By actively engaging the user, the system functions to distill fragmented information, effectively narrowing down broad, abstract concepts into precise, mechanistically grounded, and testable hypotheses. By keeping the human expert in the loop, PersonaAI ensures that generative creativity remains constrained by biological plausibility and clinical relevance (Figure 1A).

To move from theory to validation, the framework employs a “Hypothesis Evaluation Agent” (Figure 1B). Instead of following a fixed script, this agent dynamically coordinates specialized tools to test each hypothesis. For instance, the Single-cell Atlas MCP queries a comprehensive atlas of 25 million cells pre-processed with rigorous quality control, batch correction, and cell-type annotations, ensuring that biological noise is managed through high-quality, large-scale data integration. The agent then directs sub-tools for clustering and pathway analysis, seamlessly converting abstract hypotheses into quantitative results (See Methods).

### Validation of Hypothesis Generation Capability via Temporal Cutoff Strategy

To rigorously evaluate PersonaAI’s capacity to generate scientifically valid hypotheses rather than merely retrieving existing facts, we implemented a retrospective validation strategy using a temporal cutoff. We restricted the agent’s external knowledge to literature published prior to 2020 and tasked it with generating hypotheses for established aging hallmarks. To ensure that generated hypotheses represent novel inferences rather than reproduction of existing knowledge, each hypothesis was screened against the same pre-2020 corpus to verify the absence of prior reporting. Only hypotheses without direct precedent were retained for subsequent validation. The filtered outputs were then cross-referenced against the corpus of literature published between 2021 and 2025 to verify whether the agent autonomously constructed logic that aligns with future discoveries.

We produced 114 hypotheses across the spectrum of 12 aging hallmarks (López-Otín et al., 2023). To quantify the robustness of these predictions, the hypotheses were evaluated 5 times (Supplementary Figure 1) and categorized into three levels of reliability: ‘High Confidence’ (verified in 4 to 5 out of 5 independent runs), ‘Medium Confidence’ (verified in 1 to 3 runs), and ‘Unverified’ (0 runs). Intriguingly, we observed both verifiable and unverifiable hypotheses from the recent studies in most of the hallmarks. This suggests that the system’s logic-driven outputs are not random hallucinations but are grounded in reproducible biological reasoning.

It is also of note that there are unverified hypotheses in this screening. One illustrative case is observed in the domain of “Loss of Proteostasis” (Figure 2B). While the pre-2020 consensus largely attributed proteostasis collapse to a decline in degradation machinery (e.g., autophagy or proteasome failure), PersonaAI synthesized a hypothesis centered on production fidelity. It further delineated a specific mechanism related with the ribosome defect. It further suggested that restoring translation accuracy as the therapeutic targets (Figure 2B). We conducted a manual verification to ensure that these specific mechanisms were entirely absent from the scientific literature published prior to 2020. Interestingly, the suggested hypothesis, mechanisms, and the therapeutic targets were observed in recent study. Böttger et al. quantified the age-dependent increase in translational errors (e.g., stop codon readthrough) in vivo in 2025 (Böttger et al., 2025). Stein et al. revealed that aging induces specific ribosome pausing and collisions, providing mechanistic support consistent with the predicted impairment in 2022 (Stein et al., 2022). Furthermore, Martinez-Miguel et al. showed that genetically increasing translation accuracy (via an RPS23 mutation) can extend lifespan in 2021 (Stein et al., 2022) (Figure 2B).

**Figure 2.**
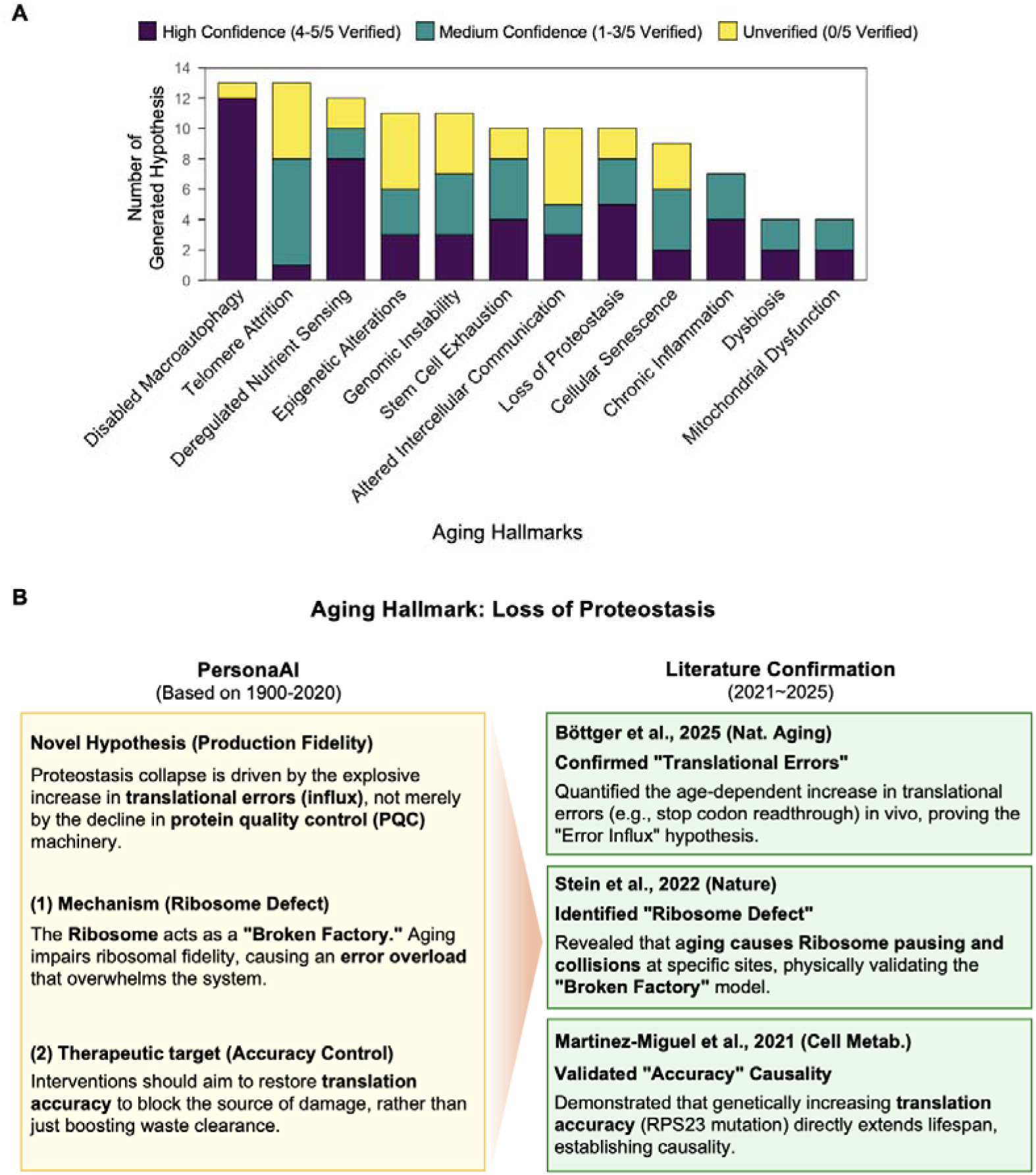
Benchmarking of PersonaAI discovery capability using a temporal cutoff strategy. (A) Distribution of 114 generated hypotheses across 12 aging hallmarks, categorized by ensemble verification confidence against 2021–2025 literature. (B) Case study of “Loss of Proteostasis” hallmark showing predicted ribosomal translational infidelity mechanisms and subsequent manual validation using post-2020 discoveries.

PersonaAI also synthesized a hypothesis that senescent cells secrete bioactive lipids that act as primary drivers of inflammation, expanding the definition of Senescence-Associated Secretory Phenotype (SASP) (Supplementary Figure 2). We also performed manual verification to ensure that it is not studied before 2020 but shown in recent studies (Byrns et al., 2024; Narzt et al., 2021; Wiley et al., 2021).

Collectively, these findings demonstrate that PersonaAI can bridge fragmented pre-2020 evidence and post-2020 findings in a way that yields testable mechanistic hypotheses. The results confirm that the framework possesses the inferential capacity to transcend simple information retrieval, establishing its utility as a powerful engine for *de novo* discovery in aging research.

### Uncovering Cellular Drivers of Aging Phenotypes: The Identification of Senescent *Cirbp*+ Hepatocytes

To evaluate the system’s capacity for cross-scale synthesis, we challenged PersonaAI to resolve the cellular context of genes previously identified only in lower-resolution studies. Specifically, we tasked the agent to identify aging-associated genes reported in bulk tissue repositories that currently lack single-cell resolution and to propose a plausible cellular mechanism for their action (Figure 3A). This discovery workflow was executed using the structured prompt logic detailed in Supplementary Table 1.

**Figure 3.**
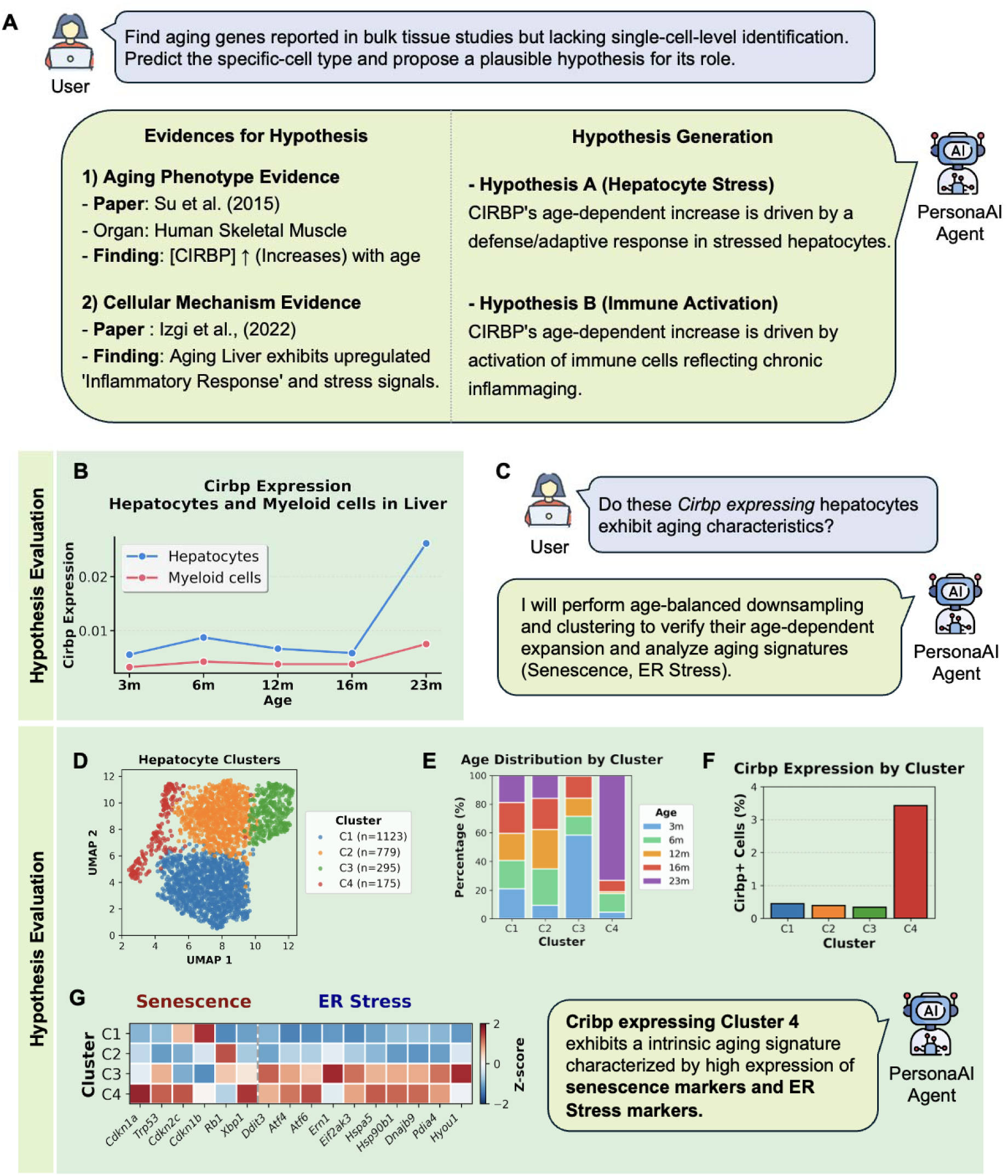
Identification of senescent Cirbp+ hepatocytes through cross-scale synthesis. (A) Discovery workflow bridging bulk tissue findings with single-cell level mechanistic identification. (B) Expression profile of Cirbp in liver hepatocyte and myeloid lineages, indicating hepatocyte-specific age-dependent upregulation. (C) Narrative of AI-human interactive inquiry regarding the aging characteristics of Cirbp-high hepatocytes. (D) UMAP visualization of hepatocyte clusters (C1–C4) derived from unsupervised clustering. (E) Cluster proportion changes across age groups (3m–23m) highlighting the selective expansion of cluster C4. (F) Relative Cirbp expression across clusters suggesting cluster C4 as a potential driver of age-associated induction. (G) Molecular signatures of senescence (e.g., Cdkn1a, Trp53) and ER stress (e.g., Hspa5, Atf4) characterizing the Cirbp-enriched cluster C4.

PersonaAI picked Cold Inducible RNA Binding Protein (CIRBP) as a target for further study using scRNAseq. To obtain it, PersonaAI linked the “Phenotypic Evidence” of systemic *CIRBP* upregulation observed in human skeletal muscle bulk transcriptomics (Su et al., 2015) with the “Mechanistic Evidence” of liver-specific stress signals reported in independent studies (Izgi et al., 2022). CIRBP is a stress-responsive RNA-binding protein that has been implicated in inflammatory signaling and reported to show age-associated regulation in multiple contexts (Barth et al., 2021; Qiang et al., 2013). Based on these evidences, PersonaAI formulated two competing hypotheses to distinguish the most plausible cellular driver mechanism. 1) Hypothesis A (Hepatocyte Stress): *CIRBP* induction represents an intrinsic defensive response of hepatocytes against oxidative or proteotoxic stress and 2) Hypothesis B (Immune Activation): the signal stems from activated resident macrophages, specifically Kupffer cells, reflecting chronic inflammaging. To further clarify the hypothesis, PersonaAI extracted the longitudinal expression profile of *Cirbp* to assess lineage-specific trends. This initial screening revealed that while myeloid populations displayed minimal changes, the hepatocyte lineage exhibited a robust, age-dependent upregulation of *Cirbp* (Wilcoxon test, *p* < 0.00001) (Figure 3B).

Because the atlas analysis indicated that age-associated Cirbp was upregulated in hepatocytes, we focused subsequent analyses on hepatocyte-intrinsic stress programs rather than immune-driven explanations: “Do these *Cirbp*-high hepatocytes exhibit specific senescence characteristics?” (Figure 3C). In response, PersonaAI downsampled the hepatocyte population to 2,372 cells, performed unsupervised clustering, and identified a distinct subpopulation associated with the highest levels of Cirbp expression (C4 in Figure 3D, 3E). Temporal distribution analysis demonstrated that cells in C4 are rarely found in young mice (3 months) but undergo a dramatic expansion during aging, becoming a dominant subpopulation by 23 months (Figure 3E, F). Finally, the agent performed deep phenotyping to characterize the molecular identity of this *Cirbp*-enriched cluster. The analysis revealed that C4 is defined by a specific intrinsic stress signature rather than general inflammation, exhibiting upregulation of canonical senescence markers (e.g., *Cdkn1a*, *Trp53*, *Cdkn2c*) and key Endoplasmic Reticulum (ER) stress regulators (e.g., *Hspa5*, *Ern1*, *Atf4*) (Figure 3G). Collectively, these results demonstrate that PersonaAI serves as a powerful discovery engine that augments the researcher by synthesizing novel findings through an integrated cycle of hypothesis generation and rigorous genomic validation. By integrating literature-based reasoning with high-dimensional single-cell analysis, the framework successfully transforms fragmented biological data into actionable, evidence-based insights.

### PersonaAI identifies the vascular niche as a driver of male-specific ASPC decline

To facilitate the strategic distillation of research inquiries, we implemented a hierarchical Meta-Prompting architecture that operates through three-step hierarchical phases to concretize the hypothesis (Method). Drawing upon the principles of Step-Back Prompting (Zheng et al., 2024), this framework compels the agent to first abstract the initial query into fundamental biological principles before converging on specific molecular targets. This top-down approach ensures that the resulting hypotheses are grounded in broad physiological contexts while maintaining the precision required for experimental validation.

This prompt (Supplementary Table 2) led us to 3 themes in sex-dependent perigonadal adipose tissue (PGAT) aging. We specifically targeted this tissue because, despite the profound sexual dimorphism well-documented in metabolic aging (Mauvais-Jarvis et al., 2020), the specific cellular determinants and niche-derived signals driving these divergent trajectories remain largely elusive even in recent transcriptomic landscapes (Emont et al., 2022).

The investigation was initiated by a broad query to “analyze key themes in sex-dependent PGAT aging.” In response, PersonaAI proposed three distinct themes. From these options, we selected “Theme 2: Adipose Stem/Progenitor Cells (ASPCs) Regenerative Decline” for further characterization, prioritizing it over the well-saturated topic of general inflammaging to maximize novelty (Figure 4A, Supplementary Table 3).

**Figure 4.**
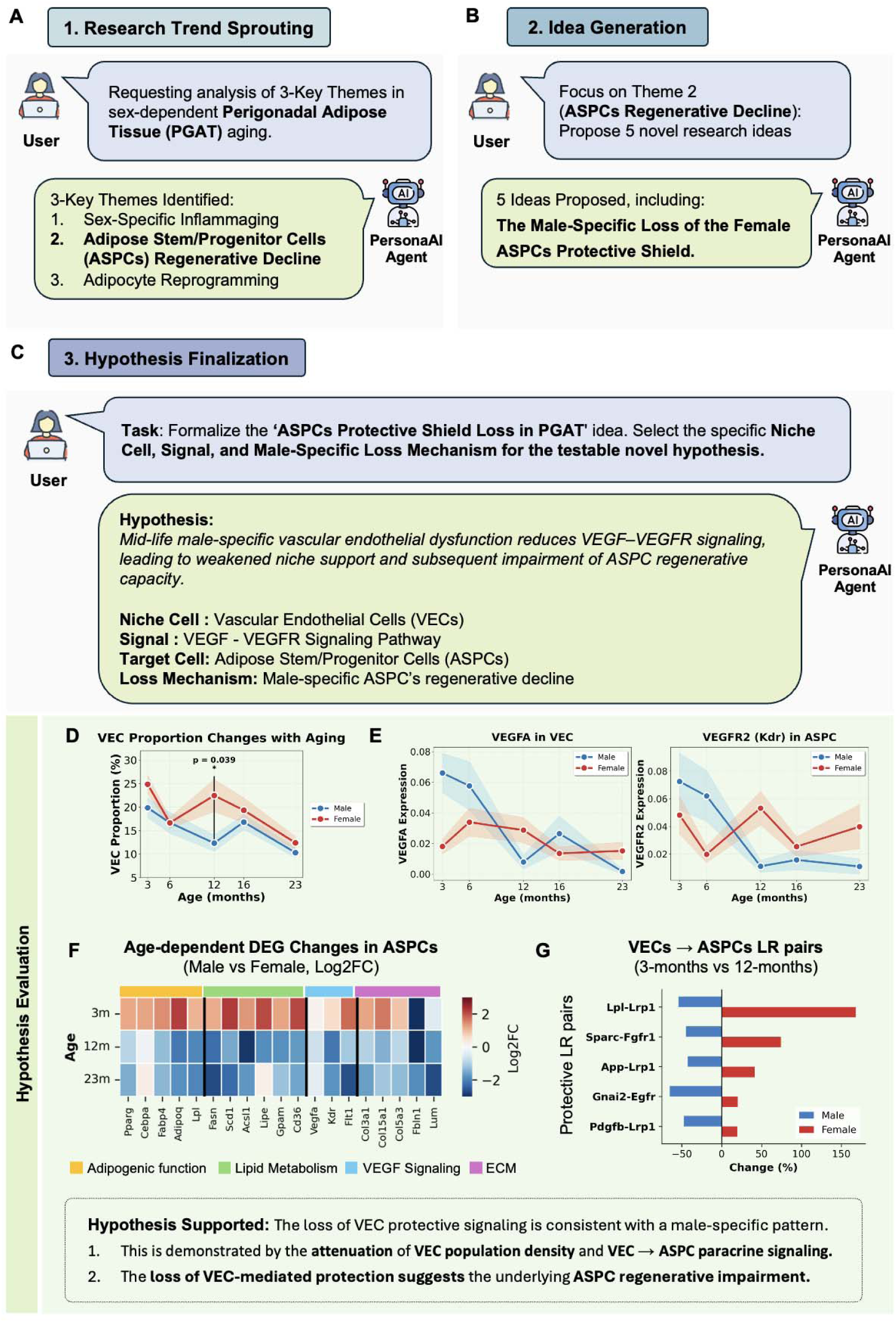
Uncovering the vascular niche as a driver of male-specific ASPC decline. (A) Research trend sprouting phase identifying three high-impact themes in sex-dependent PGAT aging. (B) Idea generation phase proposing mechanistic narratives for male-specific regenerative impairment. (C) Hypothesis finalization model integrating the VEGF-VEGFR signaling axis and niche dysfunction. (D) Age-dependent reduction in VEC population proportion specifically in the male cohort. (E) Expression levels of ligand Vegfa in VECs comparing male and female trajectories. (F) Receptor expression of Vegfr2 (Kdr) in recipient ASPCs, showing parallel decline in males. (G) Ligand-receptor (LR) interaction analysis between VECs and ASPCs indicating a male-specific loss of pro-regenerative paracrine signaling.

Subsequently, focusing on the selected theme, the agent generated five candidate research ideas. We evaluated these proposals based on biological plausibility and alignment with aging hallmarks and selected ‘Idea3: The Male-Specific Loss of the Female ASPCs Protective Shield’ as the most compelling candidate due to its strong potential connection to the “Altered Intercellular Communication” of aging hallmark (Figure 4B, Supplementary Table 4). To translate this abstract narrative into a testable mechanism, we instructed the PersonaAI to formalize the selected idea. PersonaAI structured the final hypothesis by explicitly defining the biological entities involved. The system postulated that the regenerative decline in male ASPCs is not intrinsic but is actively driven by the deterioration of the Vascular Endothelial Cell (VEC) niche and the consequent loss of the VEGF-VEGFR signaling axis (Figure 4C, Supplementary Table 5).

To validate the AI-generated hypothesis positing a male-specific collapse of the vascular niche, we executed a targeted validation pipeline on the sex-stratified PGAT single-cell atlas using PersonaAI. We observed that the proportion of Vascular Endothelial Cells (VECs) remained stable in aging females. However, aging males exhibited a significant and progressive reduction in VEC abundance (*P*=0.039) (Figure 4D). This structural decline suggests a physical loss of the supportive niche specifically in the old male mice.

Next, we interrogated the molecular integrity of the hypothesized signaling axis. Consistent with the population loss, VECs in aging males displayed a marked downregulation of the ligand Vegfa, whereas female VECs maintained robust expression levels throughout the lifespan (Figure 4E). Concurrently, the recipient ASPCs in males showed a parallel decline in the expression of the primary receptor, Kdr (VEGFR2), suggesting that VEGF signal transmission is likely reduced due to the concurrent loss of both the ligand and its receptor (Figure 4F).

Finally, to quantify the functional consequences of these alterations, the agent performed a ligand–receptor (L–R) interaction analysis between VECs and ASPCs. This analysis indicated a marked weakening of protective paracrine signaling in aging males, with several pro-regenerative and metabolic interactions—including *Lpl–Lrp1*, *Sparc–Fgfr1*, and *Pdgfb–Lrp1*—showing reduced interaction strength in aging males compared to young controls, a pattern that was notably absent in the female cohort (Figure 4G). Collectively, these computational results are consistent with the PersonaAI-derived hypothesis that male ASPC decline may be driven, at least in part, by deterioration of vascular niche support and reduced VEGF-mediated crosstalk.

To ensure that the observed sex-dimorphic patterns were not driven by technical artifacts, we performed a manual validation of the analytical pipeline. We verified that the scRNA-seq integration achieved effective batch correction while preserving biological heterogeneity across all age and sex groups (Supplementary Figure 3A-C). Furthermore, examination of the differential expression profiles confirmed that the identified signatures were statistically robust and distinct from technical noise (Supplementary Figure 4A and B). Taken together, these validation steps support the reliability of the AI-derived mechanistic model.

## Discussions

The emergence of Agentic AI marks a fundamental paradigm shift, transitioning LLMs from passive conversational interfaces to autonomous scientific engines capable of independent reasoning, strategic decision-making, and active discovery (Gibney, 2025). Unlike traditional chatbots whose function is limited to reactive information retrieval, agentic systems operate with autonomy, actively participating in the scientific process.

We engineered PersonaAI to function as an interactive cognitive co-pilot specifically optimized for the complexities of aging research. Far exceeding the capabilities of a standard knowledge retrieval tool, the system is designed to synthesize novel hypotheses by bridging disparate conceptual domains across the aging literature. As demonstrated in our validation studies (Figures 3 and 4), PersonaAI acts as a rigorous scientific partner. It engages the researcher in a dialectic process to refine theoretical models and immediately validate them against high-dimensional scRNAseq datasets.

Distinct from fully autonomous hypothesis generators (Boiko et al., 2023; Lu et al., 2024), PersonaAI is engineered with a human-centric design philosophy. In this framework, the researcher retains a pivotal role, orchestrating the scientific inquiry while leveraging the AI to accelerate learning, refine mechanistic hypotheses, and execute complex bioinformatic analyses. By functioning as an intellectual partner rather than a mere tool, PersonaAI acts as a force multiplier. It enables a single scientist to execute multi-disciplinary workflows that traditionally required a large collaborative team, effectively compressing the timeline of discovery from months to days.

PersonaAI even has a capacity to suggest a new hypothesis. A central question is if the RAG-based approach can produce truly novel hypotheses or if they are limited to summarizing and recombining existing knowledge. While critics might dismiss RAG as merely a sophisticated search mechanism (Bender et al., 2021; LeCun & Courant, 2022), advanced RAG architectures enable a powerful functional form of reasoning known as multi-hop inference (Trivedi et al., 2023). Besides, scientific novelty rarely arises from a complete scratch. It often emerges from connecting previously disparate concepts across domain boundaries. Our deployment of PersonaAI suggests that RAG-based architectures, when appropriately engineered, can indeed transcend mere information retrieval to drive genuine scientific novelty. The benchmarking we performed using temporal cut-off and the examples (Figure 2), clearly demonstrates that PersonaAI can suggest a new hypothesis.

PersonaAI derives its novelty not by inventing biological principles *de novo*, but by functioning as a hyper-efficient engine for cross-disciplinary synthesis. Leveraging a RAG architecture that indexes over 560,000 aging-related publications, the system ingests a knowledge corpus that vastly exceeds human cognitive limits, spanning siloed disciplines from evolutionary biology to clinical immunology and neuroscience. In a key demonstration of this capability, PersonaAI conceptually linked the phenotypic signature of elevated CIRBP in human skeletal muscle to the induction of hepatic (liver) inflammation (Figure 3), which has not been previously investigated. Thus, the system’s novelty emerges from its computational ability to detect latent semantic connections between disparate biological contexts that a single human specialist would be unlikely to bridge.

The capacity for novelty was demonstrated in our case study on ASPCs (Figure 4). PersonaAI did not simply report that ASPCs decline or that vasculature deteriorates. Instead, it synthesized these observations into a granular mechanistic model, proposing the specific uncoupling of the VEGF-VEGFR signaling axis as the primary driver of niche dysfunction. This moves from high-level association to a testable molecular mechanism represents a distinct value-add that goes beyond summarizing existing text.

Meta-prompting serves as the foundational governance layer of the PersonaAI architecture. This engineered prompting framework is critical for mitigating the stochasticity inherent in LLMs, ensuring that all generated outputs remain constrained within the bounds of biological plausibility. To achieve this, we developed a library of task-specific meta-prompts designed to impose rigid structural logic on the agent’s reasoning. An example is our three-phase refinement protocol, which utilizes a “Step-Back” mechanism. This architecture compels the agent to first abstract the user’s query into fundamental biological principles before progressively converging on precise, testable molecular targets (Figure 4).

Assessment using multi-agent is a crucial part in our system. By integrating agentic-AI for scRNA-seq analysis, PersonaAI can perform validation with enhanced flexibility. Rather than following a rigid script, the agent autonomously develops a customized strategy to test each biological hypothesis against the specific characteristics of the available data. Besides, it has a potential to be designed for multi-omics data. While our current focus is the biology of aging, the same agentic discovery loop could be deployed to uncover mechanisms across a wide spectrum of complex pathologies.

## Method

### Data collection and preprocessing

#### 1) Aging Atlas Repository

Our multi-agent system operates on comprehensive scRNA-seq datasets from the Mouse Aging Atlas (Sziraki et al., 2023; Zhang et al., 2025). The primary dataset comprises over 25 million cells from 17 tissue types (adipose: BAT, gWAT, iWAT; major organs: Brain, Heart, Kidney, Liver, Lung, Muscle; gastrointestinal: Stomach, Duodenum, Jejunum, Ileum, Colon; immune: B cells, T cells, Myeloid cells) spanning 5 age points from 3 to 23 months. Biological variables include sex (male/female) and strain-matched cohorts.

#### 2) Data Storage and Preprocessing

All scRNA-seq data is stored using TileDB-SOMA (Scalable Open Matrix API) (Hoffman & Wolen, 2025), a cloud-optimized columnar storage format with three components: obs (cell-level metadata: cell type, age group, sex, QC metrics), var (gene metadata: ENSEMBL IDs, gene symbols), and X (sparse expression matrix). Data is organized into experiment-level partitions by tissue type with pre-annotated cell type labels from the original atlas construction, enabling SQL-like predicate pushdown filtering and parallel I/O operations without loading entire experiments into memory. Cell filtering supports complex Boolean logic queries on cell type, age group, sex, and QC thresholds, while gene filtering operates on gene symbols rather than ENSEMBL IDs. Three sampling strategies are available: random (uniform sampling), stratified (age-balanced), and first-N (deterministic). Extracted data undergoes automated preprocessing where ENSEMBL IDs are converted to gene symbols with duplicate resolution, genes are lexicographically sorted, and output is standardized to AnnData format (h5ad) compatible with Scanpy (Wolf et al., 2018).

### Aging Hypothesis generation agent

To overcome the limitations of standard information retrieval, we designed the hypothesis generation module as a Self-Correcting RAG system driven by a LangGraph-based state machine ((Yan et al., 2024), (Ai, 2024), (Wang & Duan, 2024)). Unlike linear QA models, this system emulates the recursive reasoning process of a human researcher through an iterative “Generation-Critique-Refinement” loop.

#### 1) Knowledge Base and Retrieval

The system indexes over 560,000 aging-related publications from PubMed/PMC. To ensure high-precision retrieval, we implemented a hybrid chunking strategy and a two-stage retrieval process involving query expansion and re-ranking (Flashrank). This infrastructure supports the “Knowledge Synthesis” phase described in Figure 1A.

#### 2) Cognitive Prompt Framework (Epistemic Persona Injection)

To reduce the risk of unsupported generation, we impose explicit knowledge boundaries: hypothesis generation is restricted to pre-2021 literature, and the resulting claims are subsequently evaluated against held-out post-2021 studies and atlas-based evidence. This design is intended to mitigate hallucination risk, but it does not eliminate it; therefore, outputs are treated as hypotheses pending verification.

#### 3) Hierarchical Hypothesis Construction

The generation process adapts to the complexity of the research question using three distinct strategies, fully detailed in Supplementary Table 1-5.

##### 1. Temporal Cutoff Strategy

To evaluate the system’s predictive capability, a retrospective validation strategy was implemented by restricting the knowledge base to literature published prior to 2020. Grounded in expert personas, the agent moves beyond simple phenomenon summarization to identify fundamental upstream triggers through a “vertical linking” logic that synthesizes fragmented evidence into cohesive causal chains. Each output adheres to a four-category standard format—Hypothesis Title, Core Hypothesis, Synthetic Evidence, and Experimental Proposal. Final validation of these hypotheses was performed offline using held-out literature from 2021 to 2025 to verify their prospective accuracy

##### 2. Cross-Resolution Gap Finding

For the CIRBP case study, the agent used a “Bulk-to-Single-Cell” logic to identify cellular mechanisms underlying bulk tissue phenomena (Supplementary Table 1).

##### 3. Meta-Prompting Architecture

The hypothesis generation process follows a structured, multi-step reasoning chain—comprising Trend Scouting, Idea Generation, and Hypothesis Finalization to construct granular mechanistic models.

1. Research Trend Sprouting: The agent acts on a broad research field to analyze the landscape and identify key unexplored themes, filtering out well-saturated topics to select one high-potential area.
2. Idea Generation: Focusing on the selected theme, the agent engages in constrained brainstorming to propose novel mechanistic narratives that explain the observed phenomena.
3. Hypothesis Finalization: Finally, the agent refines the abstract idea into a structured, scientifically testable prediction, explicitly defining the Source, Signal, Target, and Mechanism.

For complex queries like sex-dimorphism, this workflow is further intensified to ensure the multi-dimensional resolution of the proposed models (Supplementary Table 2-5).

Finally, generated hypotheses undergo a self-evaluation step where the system quantitatively assesses logical completeness (0.0-1.0 scale) using an LLM-based scoring rubric (Table 1). This rubric evaluates criteria such as coverage depth, integration quality, and theoretical basis. Only hypotheses surpassing a strict threshold (0.9) are passed to the Hypothesis Evaluation Agent for in-silico validation.

**Table 1.** Evaluation Criteria for Assessing Hypothesis Relevance and Completeness.

| **Criterion** | **Rating** | **Qualitative Descriptor** |
| --- | --- | --- |
| Relevance | 0.0-0.3 | Answer largely misses most main topics or addresses them superficially |
|  | 0.4-0.6 | Answer addresses some main topics but has significant gaps in coverage |
|  | 0.7-0.8 | Answer addresses most main topics well with minor gaps |
|  | 0.9-1.0 | Answer comprehensively addresses ALL main topics with precision<br>(RESERVE FOR TRULY EXCEPTIONAL COVERAGE) |
| Completeness | 0.0-0.2 | Superficial treatment of main topics, lacking depth |
|  | 0.3-0.5 | Basic analysis of main topics with limited theoretical grounding |
|  | 0.6-0.7 | Good analysis of main topics with some theoretical foundations |
|  | 0.8-0.9 | Excellent analysis with strong theoretical grounding of main topics |
|  | 0.9-1.0 | Exceptional analysis with profound insights into all main topics<br>(RESERVE FOR WORLD-CLASS ANALYSIS ONLY) |

### Aging Hypothesis Evaluation agent

Our hypothesis evaluation system is built on the Claude Agent SDK framework, utilizing a unified intelligent agent integrated with multiple specialized MCP tool sets. This single-agent architecture orchestrates the end-to-end analytical workflow by dynamically invoking domain-specific tools for various tasks: (1) Aging Atlas MCP: TileDB-SOMA-based data extraction with SQL-like filtering and stratified sampling; (Hoffman & Wolen, 2025) (2) Single-cell Analysis MCP: Scanpy-based preprocessing, normalization, Harmony-based batch correction, and differential expression analysis, visualization; (Wolf et al., 2018) (3) Aging CCI Analysis MCP: LIANA implementation with pseudobulk DESeq2 for cell-cell interaction analysis; (Dimitrov et al., 2024) (4) Aging Trajectory MCP: Slingshot and PAGA algorithms for lineage inference; (Street et al., 2018; Wolf et al., 2019) (5) Aging Enrichment MCP: GSEApy-based gene set enrichment analysis with support for Enrichr, GSEA, and ssGSEA methods (Fang et al., 2022).

## Author Contributions Statement

B.C., G.L. contributed equally to this work. B.C. and G.L. conceived and designed the project. J.J. contributed to the development and construction of the agent framework. J.K. and T.K, designed experiments and revised the manuscript. K.W. supervised the study, obtained financial support, and revised the manuscript.

## Statistical analysis

For single-cell transcriptomic analysis, age-associated gene expression trends and differential expression were evaluated using the Wilcoxon rank-sum test. All computational procedures were executed through the PersonaAI framework, primarily utilizing Scanpy for data processing and statistical evaluation.

## Data availability

The primary single-cell RNA-sequencing (scRNA-seq) datasets used for in silico validation in this study were retrieved from the Mouse Aging Atlas. These datasets are publicly accessible via the Gene Expression Omnibus (GEO) under the accession series GSE212606 for brain data and GSE247719 for panoramic tissue data across 17 organs. The high-dimensional knowledge base used for hypothesis synthesis was constructed from over 560,000 publications indexed in PubMed and PMC.

## Suppe

**Supplementary Figure 1.**
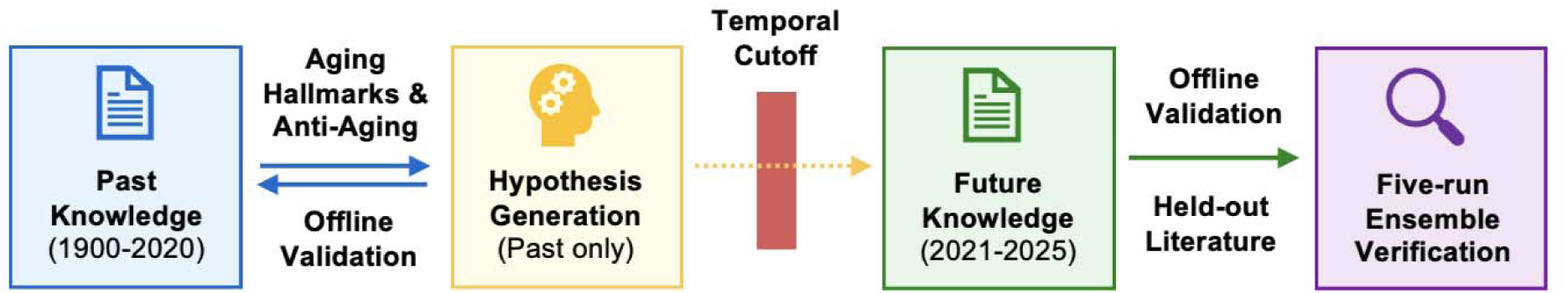
Workflow of the Temporal Cutoff Validation Strategy.

**Supplementary Figure 2.**
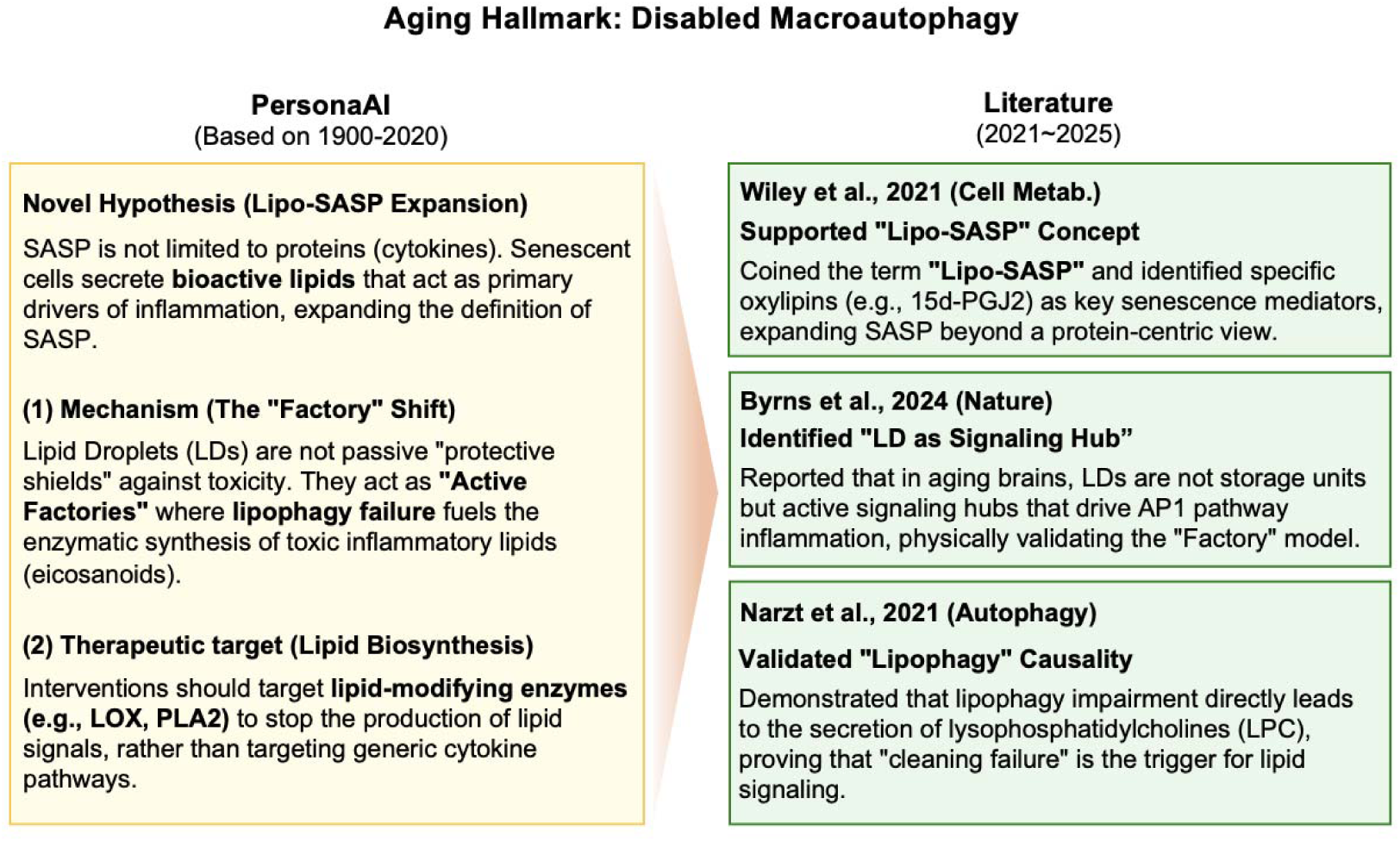
Generation and Validation of the Lipo-SASP Hypothesis.

**Supplementary Figure 3.**
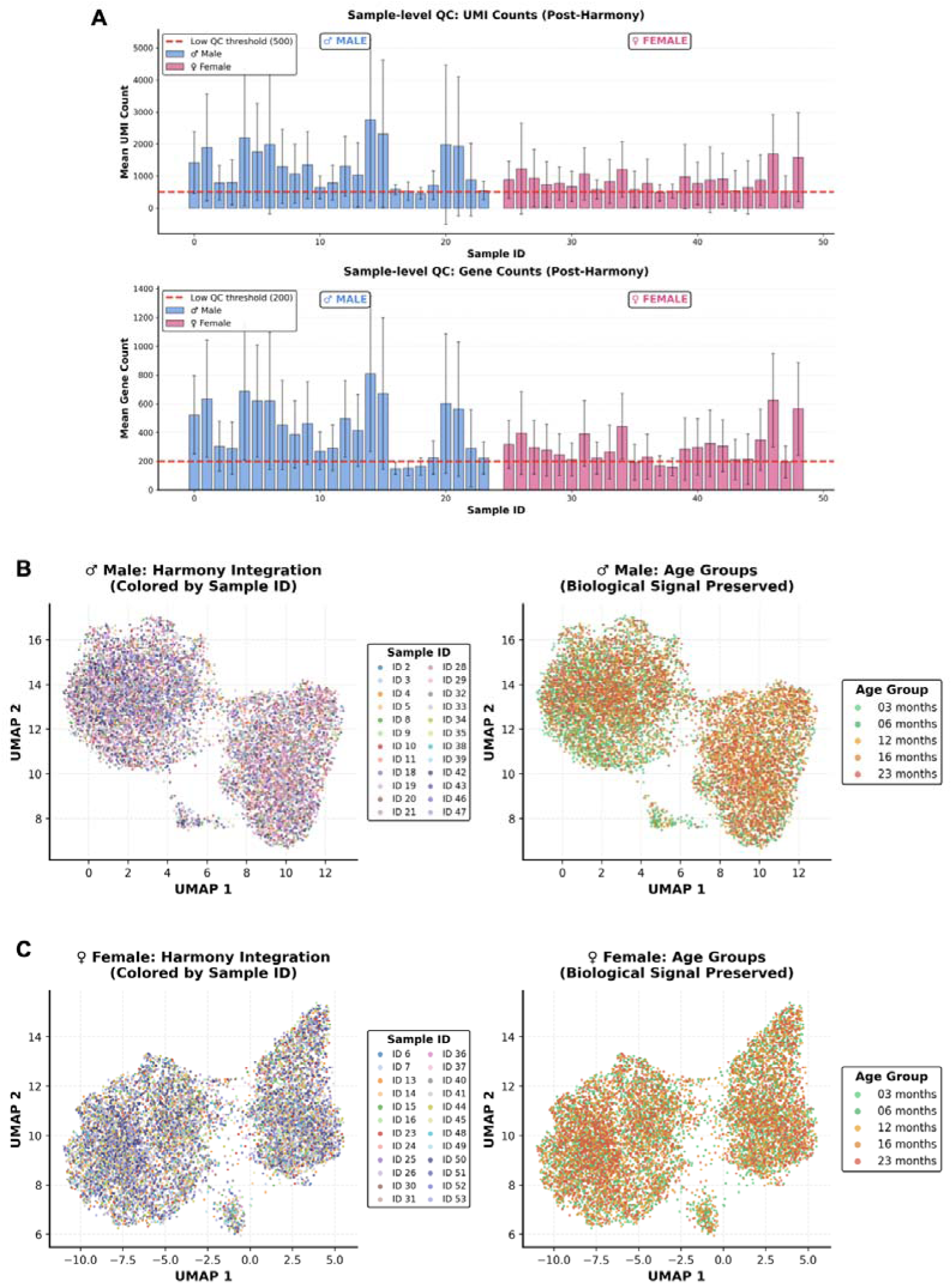
Single-cell Data Quality Control and Integration. (A) Sample-level quality control (QC) showing UMI and gene count distributions across the dataset. (B–C) UMAP visualizations for male and female cohorts demonstrating effective batch correction via Harmony and successful preservation of biological age-group signals.

**Supplementary Figure 4.**
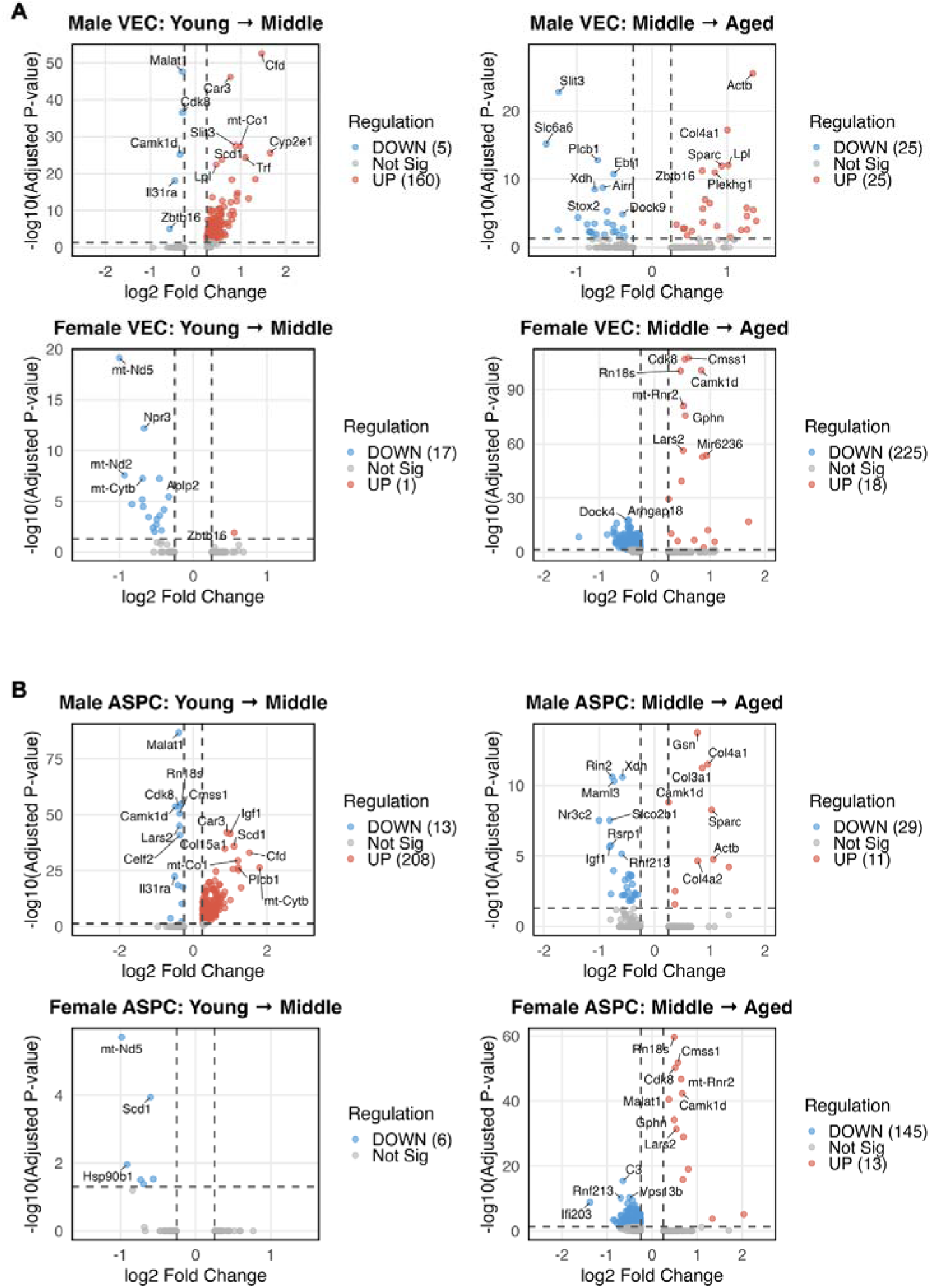
Transcriptomic Aging Signatures in VECs and ASPCs. (A) Volcano plots illustrating age-dependent differentially expressed genes (DEGs) for Vascular Endothelial Cells (VECs). (B) Volcano plots for Adipose Stem/Progenitor Cells (ASPCs). All data are stratified by sex and life stage (Young → Middle, Middle → Aged).

**Supplementary Table 1.**
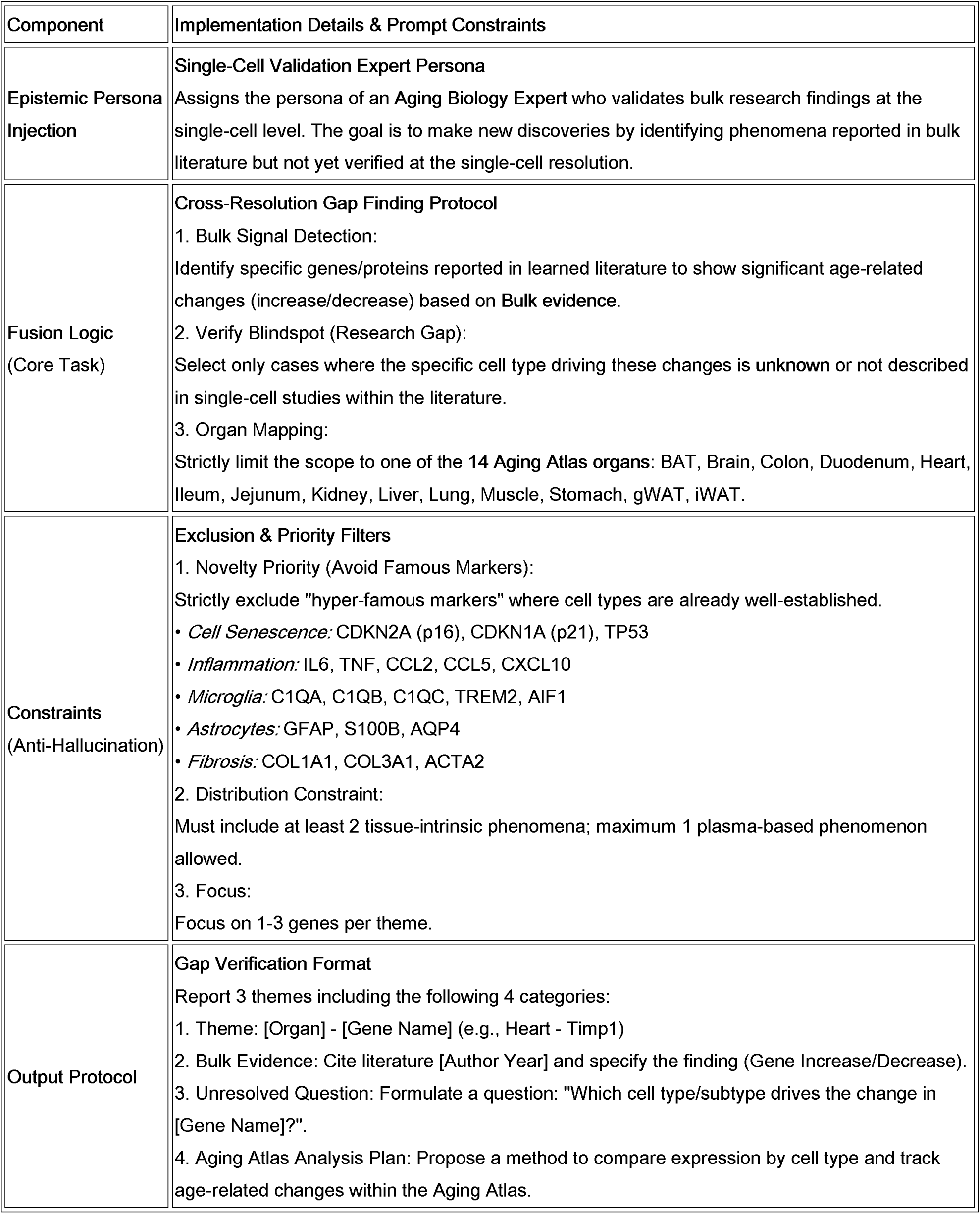
Logic Specification for Prompt concept 1 (CIRBP)

| Component | Implementation Details & Prompt Constraints |
| --- | --- |
| Epistemic Persona Injection | **Single-Cell Validation Expert Persona**<br>Assigns the persona of an Aging Biology Expert who validates bulk research findings at the single-cell level. The goal is to make new discoveries by identifying phenomena reported in bulk literature but not yet verified at the single-cell resolution. |
| Fusion Logic (Core Task) | **Cross-Resolution Gap Finding Protocol**<br>**Bulk Signal Detection:**<br>Identify specific genes/proteins reported in learned literature to show significant age-related changes (increase/decrease) based on Bulk evidence.<br>**Verify Blindspot (Research Gap):**<br>Select only cases where the specific cell type driving these changes is **unknown** or not described in single-cell studies within the literature.<br>**Organ Mapping:**<br>Strictly limit the scope to one of the 14 Aging Atlas organs: BAT, Brain, Colon, Duodenum, Heart, Ileum, Jejunum, Kidney, Liver, Lung, Muscle, Stomach, gWAT, iWAT. |
| Constraints (Anti-Hallucination) | **Exclusion & Priority Filters**<br>**Novelty Priority (Avoid Famous Markers):**<br>Strictly exclude "hyper-famous markers" where cell types are already well-established.<br>*Cell Senescence*: CDKN2A (p16), CDKN1A (p21), TP53<br>*Inflammation*: IL6, TNF, CCL2, CCL5, CXCL10<br>*Microglia*: C1QA, C1QB, C1QC, TREM2, AIF1<br>*Astrocytes*: GFAP, S100B, AQP4<br>*Fibrosis*: COL1A1, COL3A1, ACTA2<br>**Distribution Constraint:**<br>Must include at least 2 tissue-intrinsic phenomena; maximum 1 plasma-based phenomenon allowed.<br>**Focus:**<br>Focus on 1-3 genes per theme. |
| Output Protocol | **Gap Verification Format**<br>Report 3 themes including the following 4 categories:<br>Theme: [Organ] - [Gene Name] (e.g., Heart - Timp1)<br>Bulk Evidence: Cite literature [Author Year] and specify the finding (Gene Increase/Decrease).<br>Unresolved Question: Formulate a question: "Which cell type/subtype drives the change in [Gene Name]?"<br>Aging Atlas Analysis Plan: Propose a method to compare expression by cell type and track age-related changes within the Aging Atlas. |

**Supplementary Table 2.** Prompt concept 2 (Meta-Prompting/Sex-specific)

| Step | Epistemic Persona Injection | Logic Specification & Prompt Constraints |
| --- | --- | --- |
| Step 1:<br>Research<br>Trend Scouting | Sex-specific Aging Expert<br>(Focus: Tissue Heterogeneity) | **Asymmetry Detection Protocol**<br>Refuses simplistic hormone-centric explanations and identifies sexual asymmetry at the "Driver Cell" level.<br>• **Targeting:** Sets a target tissue with distinct sex-specific aging patterns (e.g., Perigonadal Adipose Tissue).<br>• **Shift Focus:** Endocrine Level → Cellular Dynamics Level.<br>• **Output:** Derives 3 verifiable asymmetric research themes for the Aging Atlas. |
| Step 2:<br>Idea Generation | Stem Cell Biologist<br>(Focus: Mechanism Divergence) | **Divergent Fate Mapping**<br>Expands the asymmetry identified in Step 1 into 5 categories of mechanistic ideas.<br>• **Anchor Logic:** Generates hypotheses centering on the "Male-Specific Loss (of Protective Signal)" vs. "Female Maintenance" pattern.<br>• **5-Category Constraints:** Forces exploration across Communication, Pathways, Patterns, Heterogeneity, and Rejuvenation to broaden the scope.<br>• **Feasibility:** Mandates the inclusion of literature hints (Indirect Evidence) and data verifiability. |
| Step 3:<br>Hypothesis<br>Finalization | Network Architect<br>(Focus: Causal Integration) | **Causal Triangulation**<br>Refines abstract ideas into verifiable Molecular Circuits to assemble the final hypothesis.<br>• **Triangulation:** Combines three elements: [Who: Niche] - [What: Ligand-Receptor] - [Why: Loss Mechanism]. |

**Supplementary Table 3.** Logic Specification applied to Prompt concept 2 (Step 1)

| Component | Implementation Details & Prompt Constraints |
| --- | --- |
| Epistemic Persona Injection | **Sex-specific Aging & Adipose Biologist**<br>The system adopts the persona of a specialist in Sex-specific Aging and Visceral Adipose Biology. It possesses a deep understanding of cellular dynamics and OSKM reprogramming, transcending the traditional endocrinological perspective to analyze tissue-level aging heterogeneity. |
| Scouting Logic (Core Task) | **Asymmetry Detection Protocol**<br>1. **Target Domain Definition:**<br>Focuses exclusively on "Perigonadal Adipose Tissue," a depot critical for metabolic disease that exhibits distinct aging trajectories between sexes (e.g., visceral accumulation patterns).<br>2. **Shift Focus (Beyond Hormones):**<br>Refuses to attribute aging differences solely to hormonal variance. Instead, it aims to identify specific "Driver Cell Populations" that actively orchestrate the aging process within the tissue context.<br>3. **Trend Analysis:**<br>Analyzes literature from the past 5 years to identify three high-impact biological themes and associated Research Gaps that have been overlooked. |
| Constraints (Context Awareness) | **Single-Cell Validation Readiness**<br>Proposed themes must not remain abstract theories but must be verifiable using Aging Atlas single-cell data (scRNA-seq). The output must highlight opportunities to elucidate "Cell-Cell Communication (Cell-Chat)" networks or distinct "Cellular Subtypes" that drive sexual dimorphism. |
| Output Protocol | **Structured Theme Report (3 Themes)**<br>Each theme must be reported with the following four mandatory components:<br>1. **Core Concept:** A summary of how aging manifests asymmetrically between sexes.<br>2. **Driver Cell Characteristics:** Identification of the specific cell types driving this asymmetry and the rationale.<br>3. **Limitations of Current Research:** A critique of existing studies, particularly the lack of rigorous sex-comparative analyses.<br>4. **Opportunities for Single-Cell Analysis:** Specific points for validation, such as unique signaling pathways or rare cell populations. |

**Supplementary Table 4.** Logic Specification applied to Prompt concept 2 (Step 2)

| Component | Implementation Details & Prompt Constraints |
| --- | --- |
| Epistemic Persona Injection | **Stem Cell & Single-Cell Biologist**<br>The system adopts the persona of an authority in single-cell transcriptomics and stem cell biology. It is tasked with going beyond descriptive observation to identify "Unexplored Territories" in sex-specific aging mechanisms, specifically focusing on the microenvironmental cues regulating stem cell exhaustion. |
| Generation Logic (Core Task) | **Divergent Fate Mapping & Mechanism Transplantation**<br>1. **Input Context (Anchor):**<br>The logic is anchored on the key finding from Step 1: *"Early depletion and hypertrophy of male ASPCs versus the proliferative maintenance of female ASPCs."*<br>2. **5-Category Ideation Strategy:**<br>The system is constrained to generate hypotheses across five mandatory categories to ensure diverse mechanistic exploration:<br>• **Cat A (Communication):** Novel cell-cell interaction pairs (e.g., Macrophage → ASPC signaling).<br>• **Cat B (Pathways):** Overlooked non-canonical pathways (vs. canonical Wnt/BMP).<br>• **Cat C (Pattern):** Communication reconfiguration patterns, specifically focusing on the "Loss of protective signals" in males.<br>• **Cat D (Heterogeneity):** Sex-dimorphic composition of ASPC subtypes.<br>• **Cat E (Rejuvenation):** Differential responsiveness to OSKM reprogramming interventions. |
| Constraints (Context Awareness) | **Feasibility Check (Aging Atlas)**<br>Proposed ideas must be grounded in reality; the system must evaluate the "Verifiability" of each idea within the Aging Atlas dataset (e.g., detectability of specific ligand-receptor pairs). It is mandatory to provide "Literature Hints" (indirect evidence) to support biological plausibility. |
| Output Protocol | **Structured Idea Proposal (5 Candidates)**<br>Each of the five generated ideas must adhere to the following format:<br>1. **Research Question:** Explicitly defines *Who* (Sex), *What* (ASPC/Signal), and *How* (Mechanism).<br>2. **Why it matters:** Articulates the impact and novelty compared to existing literature.<br>3. **Literature Hints:** Cites indirect evidence supporting the hypothesis.<br>4. **Aging Atlas Feasibility:** Assesses validation potential (Yes/Partial) with specific markers. |

**Supplementary Table 5.** Logic Specification applied to Prompt concept 2 (Step 3)

| Component | Implementation Details & Prompt Constraints |
| --- | --- |
| Epistemic Persona Injection | **Network Architect & Verification Expert**<br>The system adopts the persona of an expert in single-cell data analysis and cell-cell communication networks. It is responsible for translating abstract ideas into concrete, verifiable molecular hypotheses. |
| Finalization Logic (Core Task) | **Causal Triangulation & Integration Protocol**<br>1. Input Context:<br>Based on the "Male-Specific Loss of Female Protective Signals" paradigm selected in Step 2.<br>2. Triangulation (3-Axis Exploration):<br>The system independently identifies three core components:<br>• **Who (Niche):** Top 3 candidate cell populations (e.g., immune or vascular cells) that potentially send protective signals to ASCs.<br>• **What (Signal):** Specific Ligand-Receptor pairs mediating the support function (Proliferation/Anti-aging).<br>• **Why (Mechanism):** The biological reason for signal termination in males (e.g., Inflammatory inhibition, Niche senescence).<br>3. Integration Strategy:<br>The system synthesizes the optimal combination into a unified hypothesis using the following formula:<br>$[Niche_X] \xrightarrow{Signal_Y} [ASC_{Function}]$<br>where $\begin{cases} \text{Male: Loss via } Mechanism_M \\ \text{Female: Maintained (Protective)} \end{cases}$ |
| Constraints (Verification) | **Concrete Verifiability (Aging Atlas)**<br>The hypothesis must be actionable. It requires explicit validation points using Aging Atlas data, specifically predicting differential expression patterns of identified markers. |
| Output Protocol | **Integrated Hypothesis Report**<br>The output must follow a strict format:<br>1. **Candidate Pool:** A list of candidates for Niche cells, Signaling pathways, and Loss mechanisms.<br>2. **Final Hypothesis Statement:** A declarative statement explaining that "Signal Y from Niche X supports ASC Function Z, but is lost in males due to Mechanism M".<br>3. **Verification Points:** Three specific predictions verifiable via the Aging Atlas (e.g., "Check if Receptor R is downregulated in Male ASCs"). |

